# Comprehensive mapping of the alternative polyadenylation site usage and its dynamics at single cell resolution

**DOI:** 10.1101/2021.12.02.471022

**Authors:** Junliang Wang, Wei Chen, Wenhong Hou, Ni Hong, Hanbing Zhong, Ting Ni, Yuanming Qi, Wenfei Jin

## Abstract

Alternative polyadenylation (APA) plays an important role in post-transcriptional gene regulation such as transcript stability and translation efficiency. However, our knowledge about APA dynamics at single cell level is largely unexplored. Here we developed single cell polyadenylation sequencing (scPolyA-seq), a strand-specific approach for sequencing 3’ end of transcripts, to investigate the landscape of APA at single cell level. By analyzing several cell lines, we found many genes using multiple polyA sites in bulk data are prone to use only one polyA site in each single cell. Interestingly, cell cycle was significantly enriched in genes showing high variation of polyA site usages. We further identified 414 genes showing polyA site usage switch after cell synchronization. Genes showing cell cycle associated polyA site usage switch were grouped into 6 clusters, with cell phase specific functional categories enriched in each cluster. Furthermore, scPolyA-seq could facilitate study of APA in various biological processes.

## Introduction

Polyadenylation is essential for eukaryotic mRNA maturation, while most genes contain multiple polyA sites(Derti et al. 2012; Wang et al. 2018b). The process of a gene using different polyA sites for generating different transcripts was called alternative polyadenylation (APA) APA resulted in many transcripts with varying 3’ends, which expanded the diversity of gene products derived from a single gene. The advances of next generation sequencing resulted in the genome-wide mapping of polyA sites, which further revealed the features of polyA sites and the regulation of APA. APA involves in many post-transcriptional regulations, such as mRNA maturation, mRNA stability, cellular RNA decay and RNA’s cellular localization(Derti et al. 2012; Lianoglou et al. 2013; Gruber et al. 2016; Wang et al. 2018b; Gruber and Zavolan 2019). APA plays an important role in cell proliferation(Sandberg et al. 2008), development (Ji et al. 2009; Lianoglou et al. 2013), neural function (Miura et al. 2013), immune response(Jia et al. 2017) and aging(Shen et al. 2019). Furthermore, recent studies found tumor cells switched polyA site usage comparing with the normal cell in various cancers (Mayr and Bartel 2009; Xia et al. 2014; Park et al. 2018; Ye et al. 2019; Wang et al. 2020), particularly cancer cells prefer to use proximal polyA site to avoid microRNA-mediated repression (Xia et al. 2014).

On the other hand, cell-to-cell variations are crucial to various biological processes including embryonic development, cell homeostasis and tissue functions (Jin et al. 2015; Ren et al. 2017; Lai et al. 2018; Wang et al. 2019; Qin et al. 2021). It is reported that even seemly homogeneous cell population displays cell-to-cell heterogeneity in response to environmental stimulation (Shalek et al. 2014). Single cell RNA-sequencing (scRNA-seq), providing an unprecedented resolution to explore cell-to-cell variation of gene expression, has become an ideal approach for identifying cellular heterogeneity, searching novel cell types, exploring developmental process, inferring regulators underlying various processes and investigating tissue micro-environment (Macosko et al. 2015; Paul et al. 2015; Tirosh et al. 2016; Yu et al. 2016; Sebe-Pedros et al. 2018; Wang et al. 2019; Qin et al. 2021). The scRNA-seq focus on quantification of RNA expression at single cell level (Jaitin et al. 2014; Klein et al. 2015; Macosko et al. 2015; Qin et al. 2021), while transcript isoforms including APA have been ignored in most studies. Although several studies investigated polyA sites at single cell level(Velten et al. 2015; Shulman and Elkon 2019; Hu et al. 2020; Ye et al. 2020), they either only analyzed the cell type specific APA(Shulman and Elkon 2019) or characterized polyA site usage(Hu et al. 2020). Our knowledge about ployA site usage and dynamics at single cell level is largely unexplored partially because the limitation of data quality.

To enhance our understanding of APA at single cell level, we developed single cell polyadenylation sequencing (scPolyA-seq), a strand-specific approach for sequencing 3’ end of transcripts. Combined with Fluidigm C1 HT IFC system, we conducted scPolyA-seq on cells from 3 cell lines, namely MDA-MB-468, HeLa and mouse embryonic fibroblasts (MEFs). We described the features of polyA site usage at single cell level. We further explored APA dynamics during cell cycle, and found 6 gene clusters with distinct polyA site usage dynamics. These results indicate that scPolyA-seq could accurately identify polyA sites and demonstrate the dynamics of polyA site usage thus provided biological insights on APA.

## Results

### scPolyA-seq generates high quality data

To develop scPolyA-seq, we modified the template-switching reverse transcription method (Smart-seq2) (Picelli et al. 2014) to capture and sequence the 3’ end of transcripts (Fig. 1A). In brief, primers containing polyT and cell barcode for labeling the 3’ end of transcripts were added after cell lysis. Full-length cDNA was synthesized by template-switching reverse transcription, amplification and tagmentation with Tn5 transposases. Afterwards, DNA fragments at the 3’ end of the cDNA were captured by targeted PCR, and sequencing indices were added for amplification. The libraries were sequenced, and then polyA supporting reads were mapped to the genome for polyA site identification.

**Figure 1.**
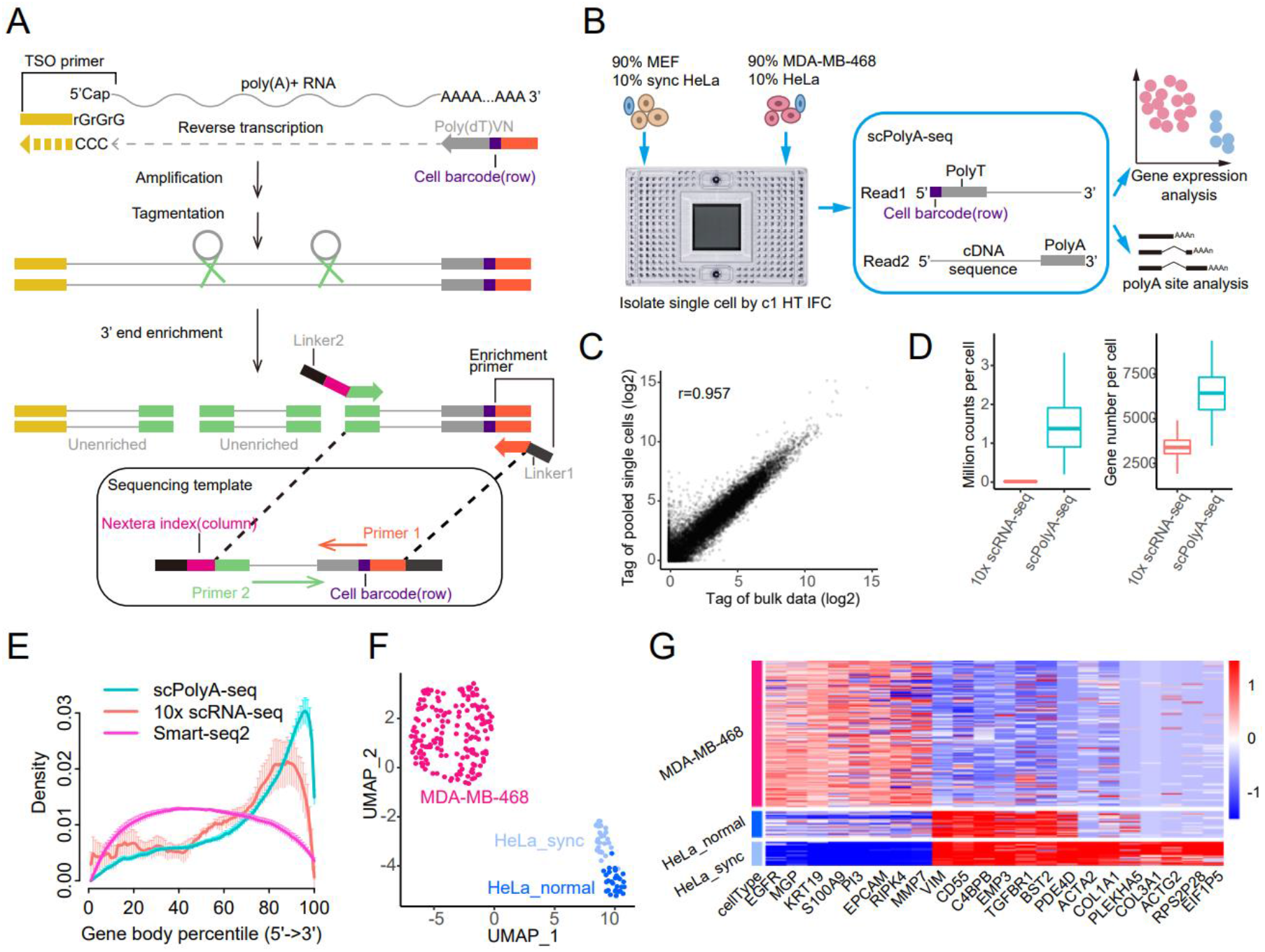
scPolyA-seq and scheme of this study. (*A*) Scheme of scPolyA-seq. The 3’end of mRNA is captured and sequenced. Read 1 carry cell barcode, while Read 2 may contain polyA sites and sequences nearby. Read 2 is used for identifying the PolyA sites and calculating gene expression. (*B*) Experiment design of this study. Cell suspensions of MDA-MB-468 and MEF (both mixed 10% HeLa cells) are loaded into two independent inlets of C1 HT integrated fluidics circuit (IFC), thus up to 400 cells can be captured for each cell suspension. (*C*) Pearson correlation of gene expression between pooled scRNA-seq and bulk RNA-seq data of MDA-MB-468. (*D*) scPolyA-seq generates much more reads and detects more genes for each cell compared with 10x scRNA-seq. (*E*) Reads from scPolyA-seq showed the sharpest peak at 3’end of each gene, compared with 10x scRNA-seq and Smart-seq2. (*F*) UMAP plot of the 3 human cell line subpopulations, namely MDA-MB-468, normal HeLa and synchronized HeLa. (*G*) Heatmap of cell subpopulation specific genes.

We designed to analyze the APA of three cell lines: triple-negative breast cancer cell line (MDA-MB-468), cervix adenocarcinoma cell line (HeLa) and mouse embryonic fibroblasts (MEFs). Fluidigm C1^™^ Single-Cell Autoprep System (Fluidigm, South San Francisco, CA, USA) was used for scPolyA-seq. Cell suspensions of MDA-MB-468 (containing ∼10% HeLa cells) and MEFs (containing ∼10% synchronized HeLa cells) were loaded into two independent inlets of a C1 HT integrated fluidics circuit (IFC), thus up to 400 cells could be captured for each cell suspension (Fig. 1B). Each well was carefully examined under a white field microscope (Supplemental Fig. S1A). The scPolyA-seq libraries were prepared and sequenced. Reads were demultiplexed to different single cells using cell barcode and mapped to reference genome (hg19 or mm10). After filtering and qualification control, 557 single cells (with 222 human cells and 335 MEFs) (Methods; Supplemental Fig. S1B,D) were left for further analysis. Two bulk libraries using small number of MDA-MB-468 were manually constructed following scPolyA-seq protocol.

The read counts between pooled scPolyA-seq data and bulk data generated are highly correlated (r=0.957), suggesting the high reproducibility of scPolyA-seq (Fig. 1C). On average, scPolyA-seq generated 1.46 million reads per cell, which is 96-fold more than that of 10x Genomics (Student’s t-test, P=1.68×10^−77^) (Fig. 1D). Furthermore, scPolyA-seq detected 6,379 expressed genes per cell on average, which is 1.87-fold more than that of 10x Genomics (Student’s t-test, P=4.33×10^−92^) (Fig. 1D). As expected, scPolyA-seq generated much less reads and detected much less genes per cell than Smart-seq2 (Supplemental Fig. S1F). Compared with 10x Genomics and Smart-seq2, the reads from scPolyA-seq were more concentrated at 3’ end of genes (Fig. 1E). MDA-MB-468, normal HeLa and synchronized HeLa showed up on UMAP (Fig. 1F; Supplemental Fig. S2A). Each cell subpopulation exhibited subpopulation specific expressed gene (Fig. 1G; Supplemental Fig. S2B).

### Identification and annotation of polyA site in single cells

To take advantages of high quality scPolyA-seq data and avoid the bias in public polyA site databases, we developed scPolyA-pipe for *de novo* identification of polyA site using scPolyA-seq data. In brief, scPolyA-pipe performs peak calling on the pooled reads from all cells for identifying the potential polyA sites. The reads on each potential polyA site in each cell were counted for further analyses (Supplemental Fig. S3A,C). Using scPolyA-pipe, we identified 20,222 polyA sites after merging adjacent polyA sites. Motif enrichment analyses of sequences around polyA sites showed hexamer motif A[U/A]UAAA and its variants significantly overrepresented at 20nt upstream of polyA sites (Fig. 2A; Supplemental Fig. S3D,E), consistent with previous studies(Lee et al. 2007; Wang et al. 2018a).

**Figure 2.**
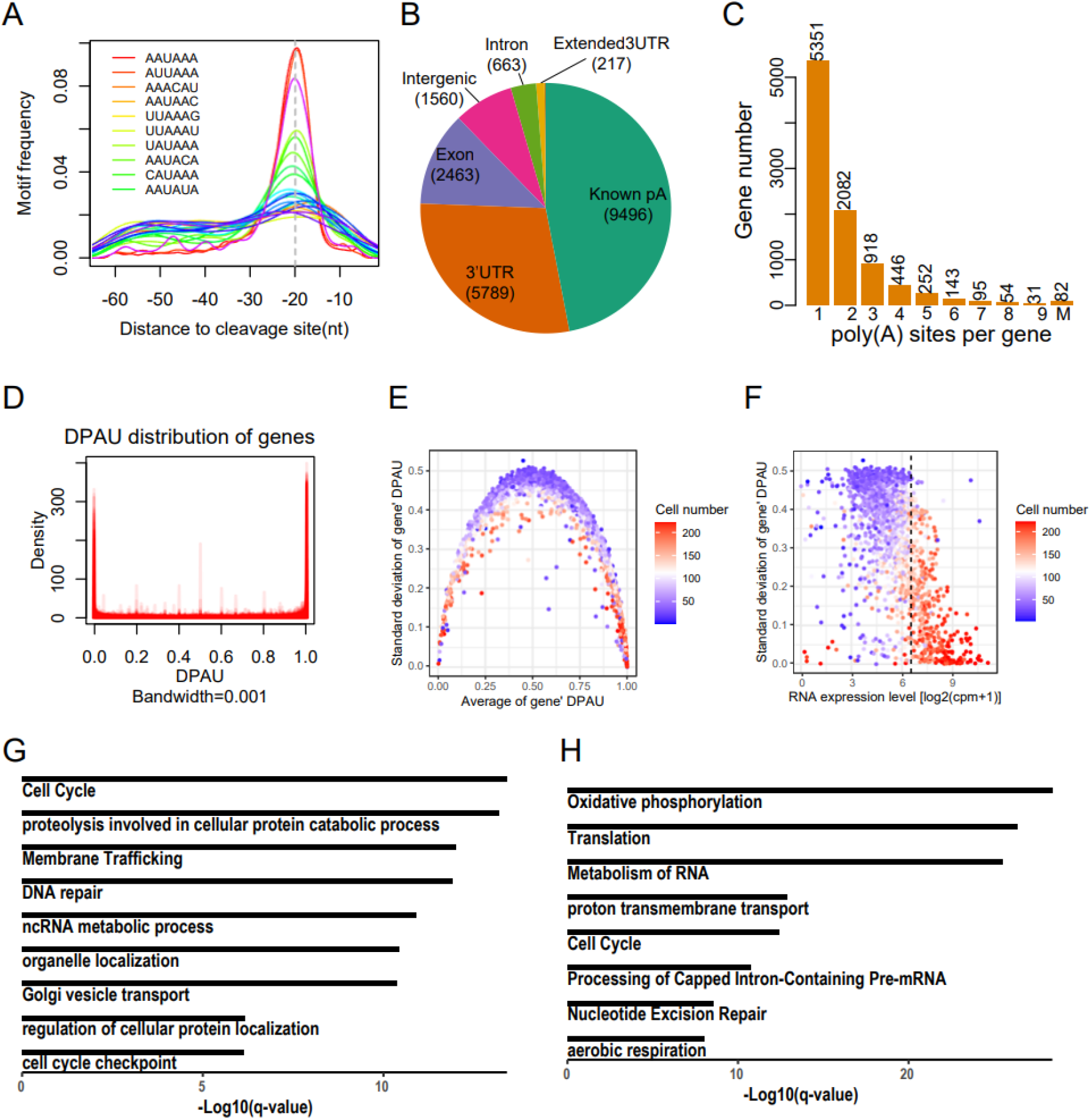
Features of polyA site usage at single cell level. (*A*) The enriched motifs near the detected PolyA sites. (*B*) Genomic location annotation of polyA sites. (*C*) Histogram of number of polyA sites per gene, with 43.4% genes contain more than one polyA sites. (*D*) Density of DPAU showing genes with 2 polyA sites across cells showed a bimodal curve, indicating these genes tend to only use one polyA site in a specific cell. (*E*) The relationship between average of DPAU and standard deviation of DPAU for genes with 2 polyA sites. One dot denotes a gene. (*F*) Scatter plot of mean expression level and standard variation of distal polyA site usage for genes with 2 polyA sites. Vertical line separates high and low expressed genes. One dot denotes a gene. (*G*) GO enrichment analyses of genes with the lowest expression level and the highest variation of polyA site usage. (*H*) GO enrichment analyses of genes with the highest expression level and the lowest variation of polyA site usage.

Most *de novo* polyA sites identified by scPolyA-seq resided in known polyA sites (46.96%), followed by 3’ UTRs (28.63%) and exon (12.18%) (Fig. 2B). More than half of these polyA sites could be found within 12nt of nearest sites in PolyA_DB2(Lee et al. 2007) (Supplemental Fig. S3F,G). These polyA sites were annotated into 9,454 genes, of which 43.3% genes contain at least two polyA sites (Fig. 2C).

### Genes using multi-polyA sites in bulk data are prone to use only one polyA site in each cell

About 43.4% genes used ≥2 polyA sites in our pooled scPolyA-seq (Fig. 2C). It is interesting to explore whether the observed multi-polyA site usage is constitutive in single cells, or simply caused by different cells using different polyA sites. We analyzed the simplest case, in which each gene has a distal polyA site and a proximal polyA site. We measured polyA site usage of each gene across these cells using **d**istal **p**oly**A** site **u**sage index (DPAU), similar to PDUI in Xia et al. (Xia et al. 2014). DPAU ranges from 0 to 1, with higher DPAU representing higher distal polyA site usage. The density of DPAU showed a prominent bimodal curve (Fig. 2D; Supplemental Fig. S4A), implying a gene tends to use only one polyA site in each cell, even these genes used multiple polyA sites in the pooled scPolyA-seq data. The genes with average DPAU near 0 or 1 showed low cell-to-cell variation on polyA site usage, while the genes with average DPAU near 0.5 showed high cellular heterogeneity on polyA site usage (Fig. 2E). We defined the phenomenon that reads on one polyA site accounting >95% of the total reads on this gene as mono-polyA site usage in a cell. We found majority of genes showing high fraction of mono-polyA site usage ratio (Supplemental Fig. S4B). The most significantly enriched GO terms in genes using both polyA sites in one single cell were protein localization to endoplasmic reticulum (P=7.41×10^−4^) and endomembrane system organization (P=2.57×10^−3^) (Supplemental Fig. S4C,D), potentially indicating these genes using APA to determine different location of transcripts.

### Variations of APAs and expression level are negatively correlated

We further found the variations of DPAU are negatively correlated with gene expression levels (Fig. 2F). The genes with the highest variations of DPAU and the lowest expression are enriched in cell cycle (P = 1.74×10^−18^), proteolysis involved in cellular protein catabolic process (P = 5.75×10^−18^), membrane trafficking (P=2.34×10^−16^) and DNA repair (P =3.89×10^−16^) (Fig. 2G). The genes with the lowest variations of DPAU and the highest expression are enriched in housekeeping gene related GO terms, such as oxidative phosphorylation (P =1.66×10^−33^), translation (P =8.13×10^−31^) and metabolism of RNA (P =1.17×10^−29^) (Fig. 2H). These results indicated the housekeeping genes had high expression and low APA dynamics, while the cell cycle associated genes had low expression and high APA dynamics.

### Relationships between APA and expression level

To evaluate the relationship between expression level and polyA site usage, we indexed the polyA site usage preference of each gene by calculating general DPAU (gDPAU), and conducted correlation test between gDPAU and expression level for each gene. We identified total 817 genes showed significant correlation between gDPAU and expression level, among which 222 genes showed positive correlations and 595 genes showed negative correlations (Fig. 3A; Supplemental Table S1). The genes showing positive correlation includes *NUTF2* and *CDKN2B*, while the genes showing negative correlation includes *BLOC1S5* and *KRAS* (Fig. 3B). There are significantly more genes showing negative correlations between gDPAUs and expression levels than genes showing positive correlation (binomial test, P<10^−16^). Therefore, genes prefer to use proximal polyA site when its expression increase, potentially because usage of proximal polyA sites are more efficient than usage of distal polyA sites and/or switching to proximal polyA site may escape miRNA induced expression inhibition(Xia et al. 2014). Indeed, the expression levels of genes showing negative correlation with gDPAU are significantly higher than the genes showing positive correlation with gDPAU (Fig. 3C).

**Figure 3.**
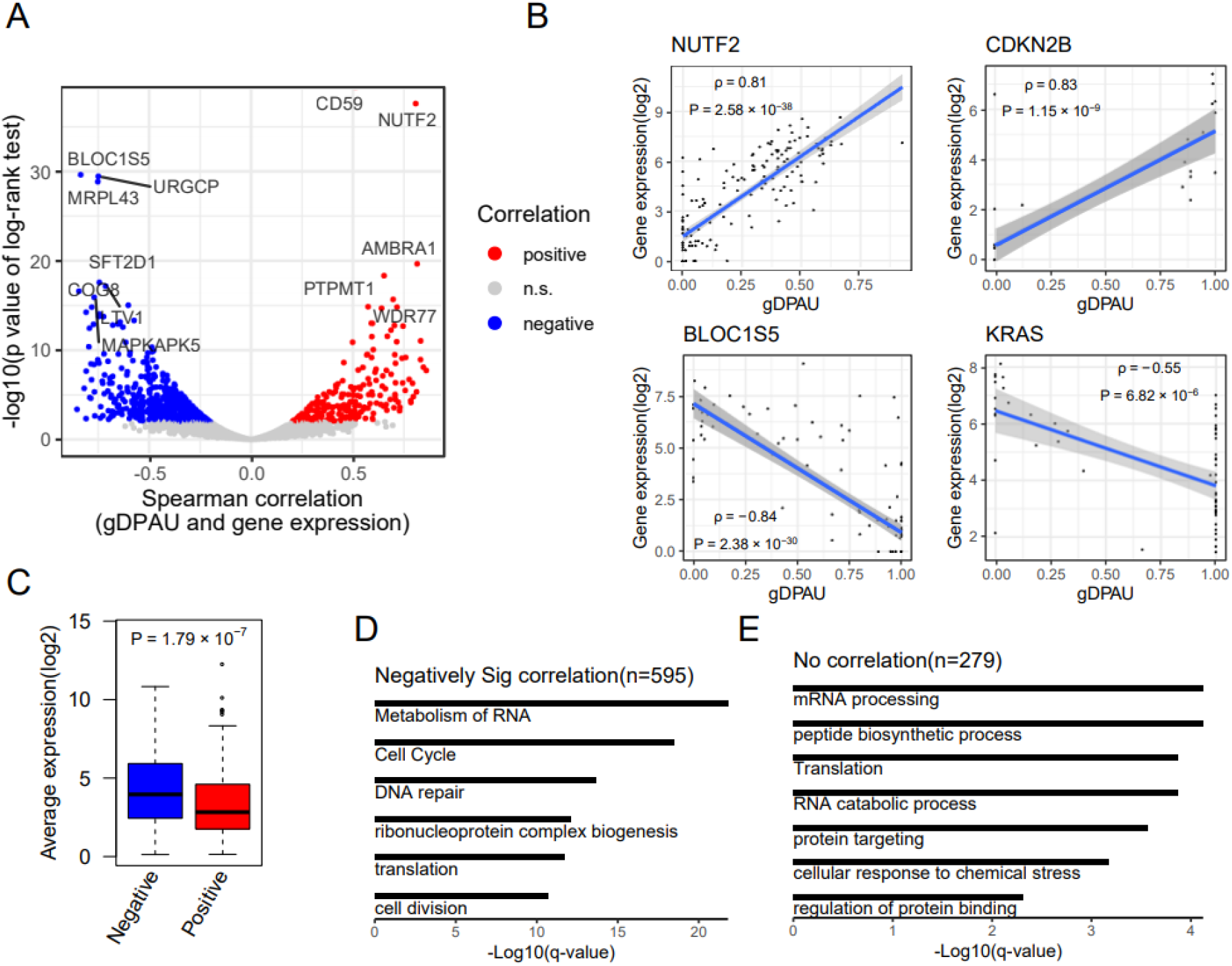
Relationship between PolyA site usage and expression level. (*A*) Identification of genes showing significant correlation between gDPAU and expression level, with each dot representing a gene. Red and blue dots represent genes with a Benjamini-Hechberg (BH) adjusted p-value of the log-rank test <0.05. (*B*) Scatter plot of gDPAU and expression for NUTF2, CDKN2B, BLOC1S5 and KRAS. Each dot represents a cell. (*C*) Expression levels of gene sets showing positive correlation and negative correlation between gDPAU and expression level. (*D*) GO enrichment analyses of genes with significantly negative correlation between gDPAU and expression (Spearman correlation<0, FDR<0.05). (*E*) GO enrichment analyses of genes showing no correlation between gDPAU and expression (|Spearman correlation|<0.02)

The genes showing negative correlations between expression levels and gDPAU were enriched in metabolism of RNA (P = 6.92×10^−27^), cell cycle (P =3.02×10^−23^) and DNA repair (P =4.68×10^−18^) (Fig. 3D), potentially indicating APA played an important role in these biological processes. The genes showing no correlation between gDPAU and expression levels were enriched in mRNA processing (P = 6.17×10^−9^), peptide biosynthetic process (P =6.61×10^−9^), translation (P =3.72×10^−8^) and RNA catabolic process (P =3.89×10^−8^) (Fig. 3E), most of which are housekeeping gene related GO terms. The enrichment of housekeeping genes further indicates housekeeping genes are unlikely affected by APA. The genes showing positive correlation between gDPAU and expression were enriched in metabolism of RNA (P = 2.33×10^−3^), Netrin-1 signaling (P = 1.53×10^−2^), RNA splicing (P = 1.53×10^−2^) and cellular responses to stress (P = 3.10×10^−2^) (Supplemental Fig. S5C). Metabolism of RNA is the most significantly enriched GO term in genes showing negative correlation and genes showing positive correlation between gDPAU and expression, which indicates that genes in metabolism of RNA showed the strongest interdependence of PolyA usage preference and expression level, either positive correlation or negative correlation.

### Changes of polyA site usage and gene expression after cell synchronization

We identified 1,019 differentially expressed genes between normal HeLa cells and synchronized HeLa cells (Fig. 4A; Supplemental Table S2), which were enriched in cellular response to growth factor stimulus (P =1.05×10^−11^), response to wounding (P =2.00×10^− 9^), response to acid chemical (P =6.61×10^−9^), cellular response to external stimulation (P =1.05×10^−7^) and response to mechanical stimulus (P = 9.77×10^−8^) (Fig. 4B). We further identified 414 genes showing significant polyA site usage switch between normal HeLa cells and synchronized HeLa cells (Fig. 4C; Supplemental Table S3), which were enriched in cell cycle associated GO terms including cell cycle mitotic (P =2.88×10^−7^) and cell cycle G2/M phase transition (P =1.41×10^−5^) (Fig. 4D). For example, *IFT20* (Fig. 4E) and *UBFD1* (Fig. 4F) are more likely to use the proximal polyA site after cell synchronization, while *GRB10* (Fig. 4G) tend to use the distal polyA site after cell synchronization.

**Figure 4.**
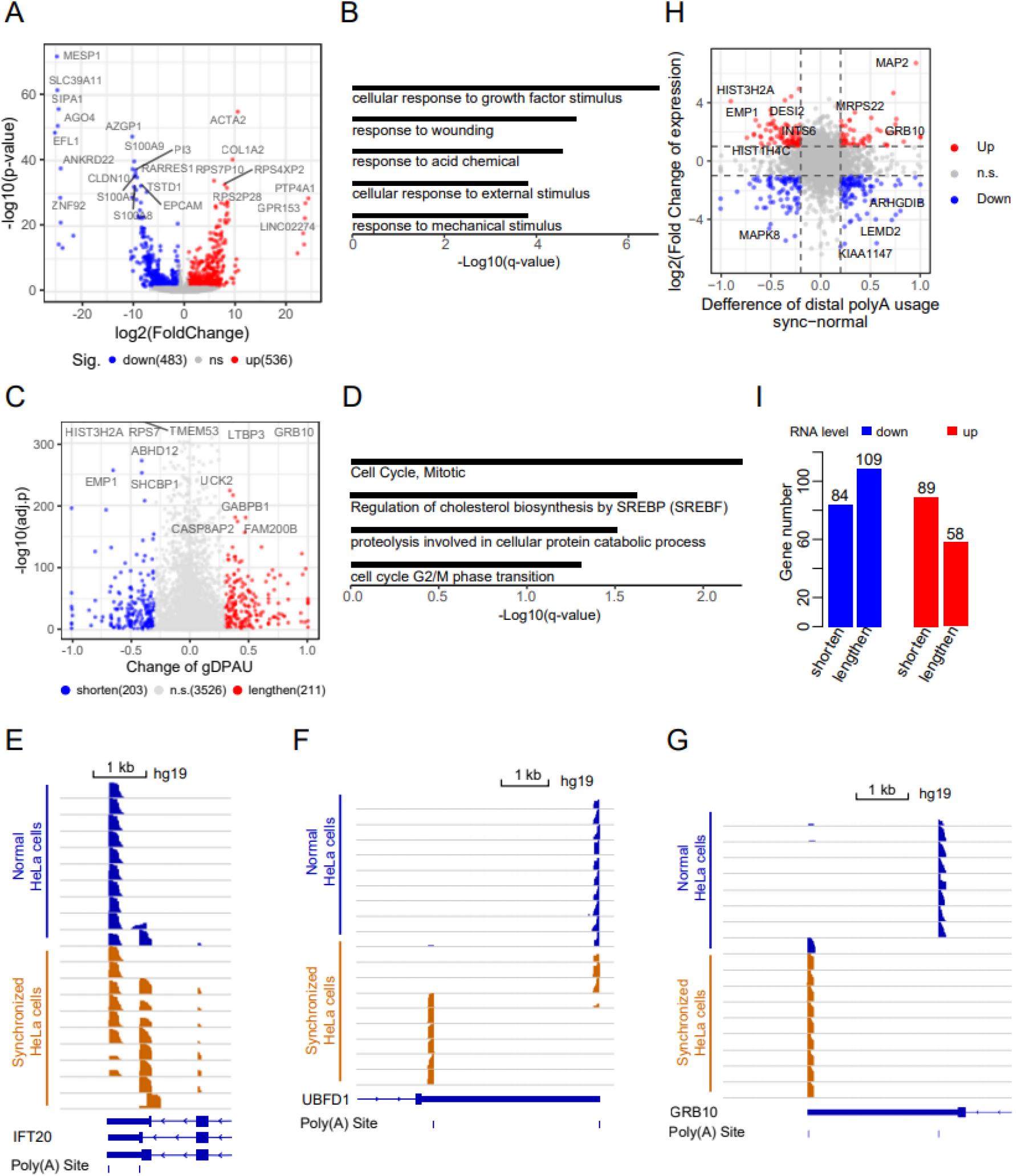
Changes of polyA site usage after cell synchronization. (*A*) Identification of differentially expressed genes between normal HeLa and synchronized HeLa by DESeq2 (FDR<0.05, log2(fold change)>1 or <-1). (*B*) GO analyses of differentially expressed genes between normal HeLa and synchronized HeLa. (*C*) Identification of genes showing significantly polyA site usage switch between normal HeLa and synchronized HeLa (FDR<0.05, χ2 test; absolute difference of gDPAU ≥0.3). (*D*) GO enrichment analyses of genes showing significant polyA site usage switch between normal HeLa cells and synchronized HeLa cells. (*E-G*) PolyA site usage changes of *IFT20* (*E*), *UBFD1* (*F*) and *GRB10* (*G*) before and after synchronization of HeLa cell. (*H*) Changes of expression level and APA level before and after cell synchronization of each gene. The fold-change of gene expression between two groups is ≥2, and the absolute difference of DPAU between two groups is ≥0.2. (*I*) Up-regulated genes and down-regulated genes are more likely to switch to proximal polyA site (P= 6.5×10^−3^; binomial test) and distal polyA site (P= 0.04; binomial test), respectively.

There is very little overlap between differentially expressed genes and polyA site usage switched genes (Supplemental Fig. S5A,B), potentially indicating regulation of expression level and regulation of APA are different mechanisms. We checked the differentially expressed genes and polyA site usage switched genes after cell synchronization in one plot to better understand their relationship (Fig. 4H). We found the up-regulated genes were more likely to use the proximal polyA sites (P= 6.5×10^−3^; binomial test) while the down-regulated genes were more likely to use the distal polyA sites (P= 0.04; binomial test). (Fig. 4I), consistent with recent studies(Xia et al. 2014; Ha et al. 2018).

### Inferring cell cycle status and cell cycle associated genes

Since we found polyA site usage switch was strongly associated with cell cycle (Fig. 4C,D), inference of cell cycle status of each cell potentially provides insights into the dynamics of polyA site usage. Each cell was assigned into a cell cycle phase using its cell cycle score according to Macosko *et al*. (Macosko et al. 2015) (Fig. 5A). The fractions of cells in cell cycle phases of synchronized HeLa are significantly different from that of normal HeLa (Χ^2^ test, P = 2.15×10^−4^). Compared with normal Hela, the number of synchronized Hela cells increased most in S phase while decreased the most in M/G1 phase (Fig. 5B). The observation is consistent with previous report that cells were arrested in S phase after double thymidine block (TdR) (Ma and Poon 2017). The expressions of *CDK4* (G1/S phase), *CCNE2* (G1/S phase and S phase), *MKI67* (G2/M phase), *CCNB2* (G2/M phase and M phase) and *CCNB1* (M phase) reached peaks at their specific representing cell cycle phase (Fig. 5C), indicating the inferred cell cycle status is reliable.

**Figure 5.**
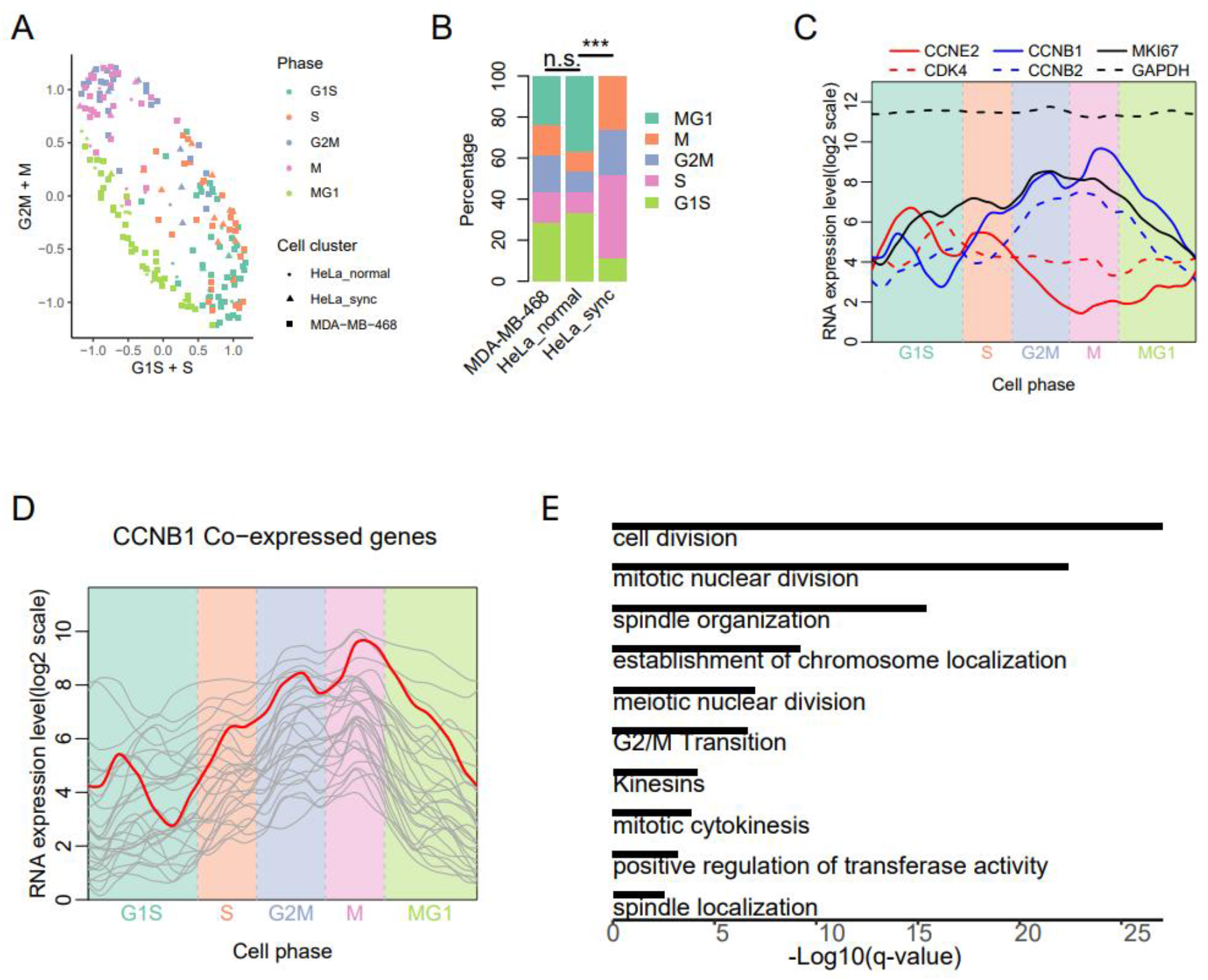
Cell cycle status and its representative genes inferred by scPolA-seq data. (*A*) Inferred cell cycle status of all cells in the 3 cell subpopulations, namely MDA-MB-468, normal HeLa and synchronized HeLa. (*B*) Bar plot of cell cycle status for the 3 cell subpopulations. The cell cycle status of synchronized HeLa is significantly different from that of normal Hela (Χ2 test, P = 2.15×10^−4^), with more cell in S phase in synchronized HeLa. (*C*) The expressions of cell cycle phase representing genes *CDK4* (G1/S phase), *CCNE2* (G1/S phase and S phase), *MKI67* (G2/M phase), *CCNB2* (G2/M phase and M phase) and *CCNB1* (M phase). Lines are smoothed using LOESS, with parameter span as 0.1. Using GAPDH as control. (*D*) Genes highly co-expressed with *CCNB1*, with Pearson correlation >0.4 and adjusted P<0.05 (n=29 genes). (*E*) GO enrichment analyses of genes co-expressed with *CCNB1*.

We identified cell cycle associated genes by inferring the co-expressed genes of each cell cycle phase representing gene (Supplemental Table S4). E.g. The Cyclin B1 (*CCNB1)* co-expressed genes showed M phase specific high expression (Fig. 5D). These genes were enriched in cell division (P =3.90×10^−32^), mitotic nuclear division (P =3.55×10^−27^), spindle organization (P =3.89×10^−19^) and establishment of chromosome localization (P =1.55×10^−12^) (Fig. 5E), which indicates the functions of these genes are M phase specific.

### Dynamics of polyA site usage switching along cell cycle

We identified the genes showing polyA site usage switches between neighboring cell cycle phases in MDA-MB-468 to explore the dynamics of polyA site usage during cell cycle (Supplemental Table S5). Venn diagrams showed majority of these genes showing polyA site usage switching were observed in one or two comparisons (Fig. 6A), which potentially indicate polyA site usage are very dynamic during cell cycle. These genes were enriched in cell cycle (P =2.89×10^−12^), regulation of cellular protein localization (P =1.05×10^−8^), and transcriptional regulation by TP53 (P =1.78×10^−7^) (Fig. 6B). We further clustered these genes using correlation of average gDPAU of each gene across 5 cell phases, resulting in 6 gene clusters (Fig. 6C). We noticed that each of the 6 gene clusters enriched specific functional GO terms (Fig. 6D). The polyA site usage dynamics of the 6 gene clusters during cell cycle were distinct from each other (Fig. 6E; Supplemental Fig. S6A). E.g., cluster1 enriched in transcription associated genes, which prefer distal polyA sites in S phase. Cluster2 enriched in regulation of dephosphorylation, which prefer distal polyA site in S phase and M phase. Cluster3 enriched in DNA repair and DNA replication, which prefer distal polyA sites in S phase. Cluster4 enriched in cell division, which prefer proximal polyA site in S phase and M phase. Cluster5 enriched in regulation of cellular protein localization, which prefer distal polyA site in M phase. Cluster6 enriched in nucleus organization, which prefer proximal polyA site in S phase. PolyA site usage switch belongs to post-transcriptional regulation that could quick response to environment stimulation, which may explain why polyA site usage switch plays an important role in cell cycle (Fig. 6D,E; Supplemental Table S6). The genes showing polyA site usage switch during cell cycle in HeLa were also grouped into 6 clusters and the gDPAU of 6 clusters showed dynamics patterns similar to MDA-MB-468 (Supplemental Fig. S6B), which indicate that polyA usage switch during cell cycle are conserved between different cell lines/types.

**Figure 6.**
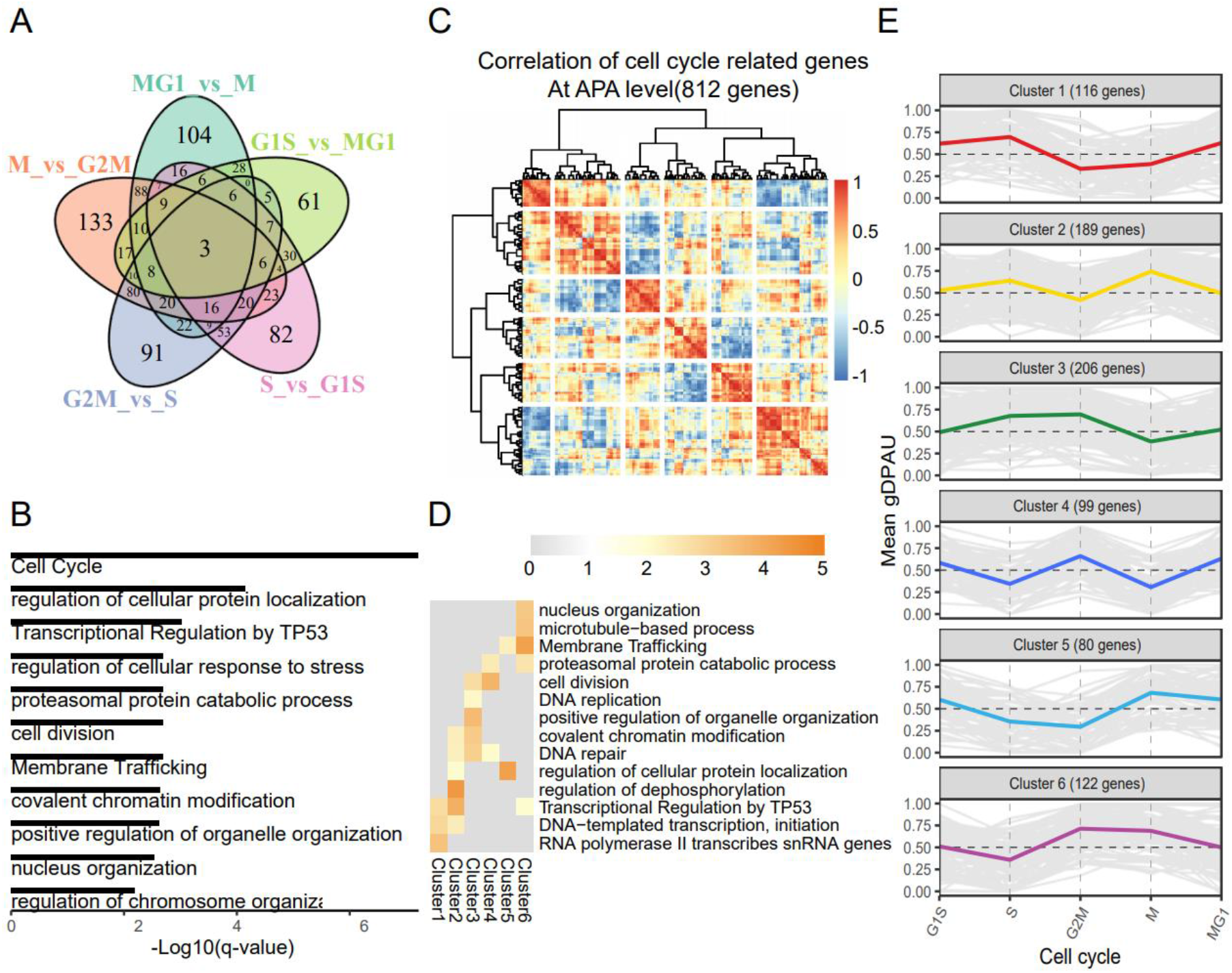
Dynamics of polyA site usage during cell cycle. (*A*) Venn diagram of genes with different polyA site usages between neighboring cell cycle phases. (*B*) GO enrichment analyses of all genes showing polyA site usage switch between neighboring cell cycle phases. (*C*) Heatmap of correlation of gDPAU of each gene across 5 cell phases, resulting genes clustered into 6 gene clusters. (*D*) GO enrichment analyses of each gene clusters. (*E*) Dynamics of gDPAU of each gene at each phase for the 6 gene clusters. Each line denotes one gene. Y-axis denotes this gene’s gDPAU at each phase. Colored lines are averaged gDPAU of each gene cluster.

Interestingly, we found the gDPAU of cluster1 and cluster6 were negatively correlated (Supplemental Fig. S6C), indicating that transcriptional associated genes and nucleus organization associated genes showed opposite pattern of polyA site usages. In particular, cluster1 and cluster6 reached maximum gDPAU and minimum gDPAU in S phase, respectively (Fig. 6E). Furthermore, negative correlation of gDPAUs between cluster3 and cluster5 indicated that DNA repair/replication associated genes and cellular protein localization associated genes showed opposite pattern of polyA site usages; Negative correlation of gDPAUs between cluster2 and cluster4 indicated that cell division and regulation of dephosphorylation showed opposite pattern of polyA site usages (Supplemental Fig. S6C). These results indicated that different gene categories have different preferences on the polyA site usage during cell cycle. These findings provide novel polyA targets to intervene cell cycle, which may facilitate therapy for cancers and other diseases.

## Discussion

Although polyA tail has been targeted for mRNA capture, scRNA-seq only focused on quantifying gene expression while ignored polyA site usages. Particularly, almost all 3’ end scRNA-seq sequences the alternative end of targeted fragment instead of polyA end to avoid the potential sequencing error caused by polyA. Therefore, it is very difficult to directly identify the polyA site for APA analyses using conventional scRNA-seq. Different from conventional scRNA-seq, scPolyA-seq sequences much close to the 3’ end of transcripts and generates high fraction of polyA-supporting reads (Supplemental Fig. S3H). The high fraction of polyA-supporting reads greatly facilitates identification of the exact location of polyA sites in the genome, thus make the scPolyA-seq more suitable for APA analyses. Furthermore, scPolyA-seq generated about 96-fold more reads per cell than that of 10x Genomics on average, which make scPolyA-seq have enough statistical power to detect polyA sites in single cell. Overall, scPolyA-seq provides a nice approach to analyze the cellular heterogeneity of APA, and provide an opportunity for exploring the relationship between gene expression and polyA site usage.

Based on analyses of MDA-MB-468, we explored the relationship between polyA site usage and expression level for better understanding their relationship. We identified 222 genes and 595 genes showing significantly positive correlation and negative correlation between expression level and distal polyA usage, respectively, indicating highly expressed genes prefer to use the proximal polyA sites instead of distal polyA sites. Indeed, the expression levels of genes showing negative correlation with distal polyA usage are significantly higher than the genes showing positive correlation with distal polyA usage. On the other hand, usage of proximal polyA site generates short RNA, which is potentially more energy efficient and easier to escape microRNA-mediated repression. Furthermore, we found the variations of APAs and expression level are negatively correlated, potentially indicating the lowly expressed genes are likely to randomly use a polyA site while highly expressed genes are prone to use a specific polyA site. In this way, we could hypothesize the relationship between polyA usage and expression: A gene with multiple polyA sites may randomly use a polyA site when its expression is low or moderate. It began to prefer one of these polyA sites when its expression increases. For these extremely high expressed genes, a gene is more likely to use the proximal polyA site than the distal site for more efficient expression.

Potentially because technology limitation for detecting polyA sites dynamics at single cell resolution, there is no systematic study on APA dynamics during cell cycle so far. After we found cell cycle is the most significantly enriched GO term in changed polyA sites usage between normal HeLa and synchronized HeLa, we realized that polyA site usage might be highly dynamic during cell cycle. We analyzed the genes showing polyA sites usage switch between neighboring cell cycle phases and identified 6 gene clusters showing distinct APA dynamic patterns during cell cycle. PolyA site usage switch belongs to post-transcriptional regulation that could quick response to environment stimulation, which may explain why polyA site usage switch plays an important role in cell cycle. Identification of the 6 gene clusters with specific APA dynamics pattern significantly improved our understanding of APA dynamics. These findings provide novel polyA targets to intervene cell cycle, which may facilitate therapy for cancers and other diseases.

## Materials and Methods

### Cell culture and cell synchronization by double thymidine block

MDA-MB-468 and HeLa cell lines were obtained from American Type Culture Collection (ATCC) and cultured in DMEM (Gibco) with 10% FBS (Gibco) in 5% CO_2_ at 37°C. MEF was prepared following our recent study(Chen et al. 2020). Synchronization of HeLa cells by double thymidine block was conducted following protocol in *Ma and Poon* (Ma and Poon 2017). In Brief, Hela cells were separated and cultured in two 25 cm^2^ flasks, for control and cell synchronization, respectively. To synchronize the cells, thymidine (2 mM, Sigma) was added to a flask and the cells were incubated for another 18h, washed 3 times with PBS, and released into thymidine-free complete medium for 9h. Thymidine (2 mM) was then added again and cultured for an additional 15 h. Hela cells in control flask were cultivated in complete medium without adding thymidine.

### scPolyA-seq

We used Fluidigm C1™ Single-Cell Auto Prep System (Fluidigm, South San Francisco, CA, USA) for scPolyA-seq library preparation. Library construction followed Fluidigm manual and protocol (DeLaughter 2018), except these steps adapted to scPolyA-seq. In brief, MDA-MB-468, MEF and HeLa cells were collected and counted in Cellometer Mini (Nexcelom Bioscience). Then cells were mixed and re-suspended to 400 cells/μL cell suspensions. Fluidigm high-throughput integrated fluidic circuit (HT IFC) with 800 wells was used for cell capture. Cell suspensions of MEF (mixed with 10% synchronized HeLa cells) and MDA-MB-468 (mixed with 10% HeLa cells) were loaded into two independent inlets of IFC (Fig. 1B), up to 400 cells could be captured for each cell suspension. HT IFC was checked under white field microscope to record the cell status in each well: empty, single, debris or dual (Supplemental Fig. S1A).

The C1 Single-Cell mRNA Seq HT Reagent Kit was used for cDNA synthesis. We started scPolyA-seq library preparation by cell lysis. Then we added primers containing polyT and cell barcode for labeling the 3’ end of transcripts. Full-length cDNA was synthesized by template-switching reverse transcription, following by amplification, and tagmentation with Tn5 transposases. Nextera XT DNA Library Preparation Kit (Ilumina) was applied to construct scPolyA-seq library. DNA fragments at the 3’ end of the cDNA were captured by targeted PCR, and sequencing indexes were added for amplification. The quality of library was examined by Qubit 3.0 Fluorometer and Agilent 2100 Bioanalyzer. Libraries separated by columns were pooled together and sequenced on Illumina HiSeq2500 to obtain 150-bp paired-end reads.

### scPolyA-seq data preprocessing and quality control

The scPolyA-seq data were demultiplexed to single cells using a script provided by Fluidigm (mRNASeqHT_demultiplex.pl). The data quality was checked by FastQC v0.11.7 (https://www.bioinformatics.babraham.ac.uk/projects/fastqc). The number of reads for each cell was essentially consistent with the cell status in each well based on microscopy check of IFC (Supplemental Fig. S1A,B). Cells that have <3.2× 10^5^ reads were filtered out (Supplemental Fig. S1C). Reads were filtered and trimmed using Trim Galore version 0.6.1 (https://www.bioinformatics.babraham.ac.uk/projects/trim_galore/). Reads from each cell were mapped to human reference genome (hg19) by STAR version 2.5.2b(Dobin et al. 2013). Gene expression level of the coding genes from GENCODE v30(Frankish et al. 2019) was quantified by htseq-count(Anders et al. 2015).

The scPolyA-seq library was strand-specific (Supplemental Fig. S1E), which may avoid some ambiguity in overlapping regions on the genome when identifying polyA sites. To remove potential outliers, we calculated the median absolute deviation (MAD) of numbers of mapped reads and expressed genes of all cells. The cells were filter out if the read counts or number of expressed genes >medians+3×MAD or <medians-3×MAD. Cells were removed if proportion of their mitochondrial RNA were outlier. Counts per million (CPM) was calculated to quantify the gene expression level since scPolyA-seq only sequenced the 3’ end of each transcript (Fig. 1E).

### Identification of cell types and cell populations

We generated scPolyA-seq data for 4 cell populations, namely MDA-MB-468, MEF, Hela synchronized and normal Hela. MEF and synchronized HeLa on the same inlet of IFC were separated by mapping to hg19_mm10 mega reference genome using STAR (Macosko et al. 2015) and plotting uniquely mapped reads (Supplemental Fig. S1D). Synchronized HeLa cells combined with MDA-MB-468 and normal Hela cells from the other inlet of IFC were projected on uniform manifold approximation and projection (UMAP) by Seurat 3.1.5(Stuart et al. 2019; Zhou and Jin 2020) (Supplemental Fig. S2A,B). Differential gene analysis was performed by DESeq2(Love et al. 2014).

### Identification of polyA site

We developed a *de-novo* polyA site identification pipeline with Snakemake (Koster and Rahmann 2018), named scPolyA-pipe, to identify polyA site from sequence data (https://github.com/WangJL2021/scPolyA-seq). To accurately locate polyA site, we only selected polyA site supporting (PASS) read, which either contains ≥10 consecutive As or with ≥6 consecutive ‘A’s at the 3’-end, for further analyses (Supplemental Fig. S3A,B). The upstream of PASS read immediately before consecutive ‘A’s is the potential sequence before polyA site and was cleaved for mapping to human reference genome (hg19). The uniquely mapped reads from each cell were saved in a bam file. All the bam files were merged into one file, in which the 3’-end of each read was consider as the polyA cleavage position. These polyA cleavage positions were used for inferring the polyA site, similar to Hoque et al. (Hoque et al. 2013). In brief, these polyA cleavage positions were merged into one polyA site cluster if they were on the same strand and within 24nt with each other. If a cluster extends >24nt, the site with the highest number of cleavage positions was identified and other sites located further than 12nt from this site were re-clustered. This process was repeated until all polyA site clusters expands <24nt. We got a total of 740,973 polyA sites/clusters without consider the mitochondrial DNA. To eliminate internal priming effect, we removed polyA sites with 6 consecutive As or more than 15 As in 20nt downstream region of the cleavage site on genome. We only kept the polyA sites supported by at least 10% percent of cells. Finally, 20,222 polyA sites passed the quality control and were used for further analyses.

### Features of identified PolyA sites

Sequences of 100nt upstream and downstream of polyA sites were extracted and base frequency was plotted (Supplemental Fig. S3E). In order to identify the enriched motifs, the 60nt upstream sequences of each polyA sites extracted by *bedtools getfasta* (Quinlan 2014) were summited to MEME (Bailey et al. 2015) to discover motifs (Supplemental Fig. S3D).

### PolyA site annotation

PolyA sites were annotated according to hg19 GENCODE v30. We annotated polyA sites located in ± 10bp of 3’ end of transcripts as known polyA. PolyA sites located in exon, intron, 3’ UTR and downstream 1kb of 3′ UTR were annotated as exon, intron and UTR3 and Extended3UTR, respectively. For polyA site falling into multiple categories, we set priority as Known pA > 3’UTR > Extended3UTR > Exon > Intron >Intergenic.

### Distal polyA site usage index (DPAU) and generalized DPAU (gDPAU)

For genes with two polyA sites, we used DPAU to quantify the percentage of distal polyA site usage for each gene in each cell. E.g., in cell *i*, the DPAU of gene *g* can be calculated as:

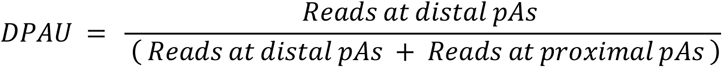

DPAU ranges from 0 to 1. In each cell, if a gene’s total supporting reads at both sites are less than 5, this gene’s DPAU in this cell will be regarded as noise and noted as NA. Otherwise, this cell will be considered as a valid observation.

For genes with more than two polyA sites, **g**eneralized DPAU (gDPAU) is used to quantify the trend of distal polyA site usage for each gene. It’s a location index weighted sum of read count percentage of gene’s each polyA sites. E.g., for a gene with n (n≥2) polyA sites, p1, p2, …, pn represents the percentages of its polyA site usage at each site from 5’-end to 3’-end.

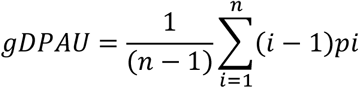

When n=2, gDPAU=DPAU. In each cell, if a gene’s total supporting reads at all sites are less than 5, this gene’s gDPAU in this cell will be regarded as noise and noted as NA. Otherwise, this cell will be considered as a valid observation.

### Identifying changes of APA between groups

Chi-squared test is used to compare the significance of gDPAU changes between groups by stacking counts at each polyA site within one group. Benjamini-Hoschberg was used for multiple comparison corrections. If a gene’s absolute mean difference of gDPAU is >0.3 and adjusted p value is < 0.01 between two groups, the gene’s APA change between two groups will be deemed as significant.

### Visualization and enrichment analyses

Reads, polyA sites and gene across cells are visualized by IGV 2.8.2(Thorvaldsdottir et al. 2013). GO representative analysis is done by Metascape (Zhou et al. 2019). False discovery rate (FDR) correction is used for multiple comparison correction.

### Cell cycle phase assignment

Cell cycle phase was assigned following a method descripted in Drop-seq(Macosko et al. 2015). Briefly, cell cycle related genes of each phase(Whitfield et al. 2002) were selected and average expression level of these gene sets were calculated. Genes in each set whose correlation with the average expression level of this set across all cells larger than 0.3 were retained. The average of gene set expression level of each phase was deemed as the cell cycle score of this phase. The cell’s phase would be assigned with the top phase score. And cells were ordered according to their phase score of each phase.

### Statistical analysis

The statistical tests and plots were conducted using R version 3.6.0 (2019-04-26).

## Data availability

All raw and processed sequencing data have been submitted to the NCBI Gene Expression Omnibus (GEO; https://www.ncbi.nlm.nih.gov/geo/) under accession number GSE178264.

## Code availability

The source code and scripts for processing scPolyA-seq are maintained in the GitHub code repository: https://github.com/WangJL2021/scPolyA-seq

## Acknowledgments

This study was supported by National Key R&D Program of China (2018YFC1004500), National Natural Science Foundation of China (31771603), Shenzhen Science and Technology Program (KQTD20180411143432337, JCYJ20170412153453623, JCYJ20180504170158430), Shenzhen Innovation Committee of Science and Technology (ZDSYS20200811144002008). We thank the Center for Computational Science and Engineering in SUSTech for computational support.

## Author Contributions

W.J., and T.N. conceived the project. W.C. did the experiment. J.W. analyzed the data, with contribution from W.H and H.Z.. W.J., Y.Q. and N.H. supervised this study. W.J. and J.W. wrote the manuscript. All authors have read and approved the manuscript.

