## Supplemental for "Comprehensive mapping of the alternative polyadenylation site usage and its dynamics at single cell resolution"

**This PDF file includes:**

Figures S1 to S6
Legends for Tables S1 to S5

**Other supplementary materials for this manuscript include the following:**

Tables S1 to S6

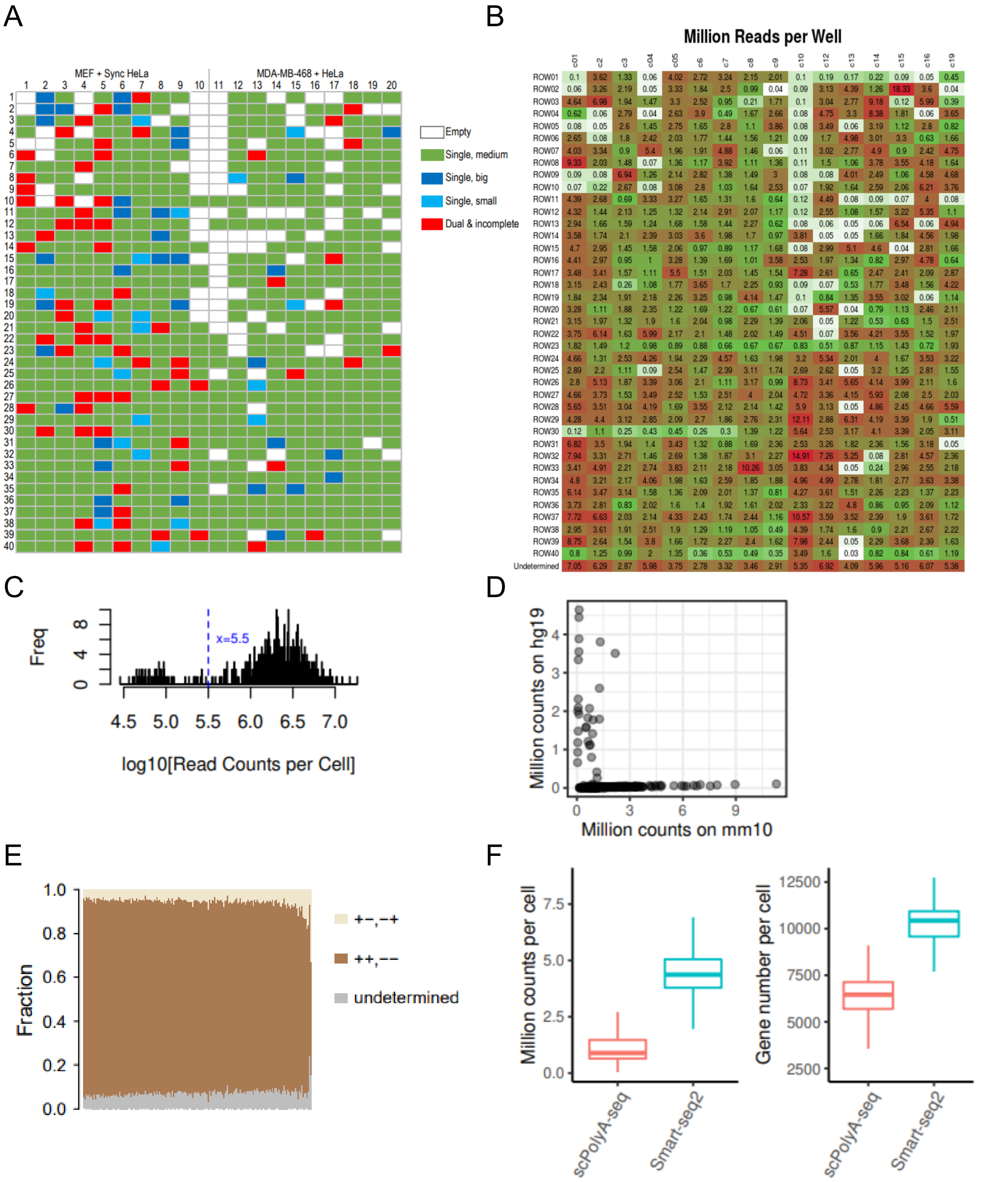


**Fig. S1. Quality control for scPolyA-seq data.** (A) Status of chip captured cells in each well under white-field microscope. (B) Heatmap of million reads per well after high throughout sequencing. (C) Histogram of counts per cell in log10 scale to determine the threshold for empty cells. (D) Uniquely mapped reads to hg19 and mm10 based on reads mapped to hg19_mm10 mega reference. One dot denotes a cell. (E) scPolyA-seq is a strand-specific library determine by RSeQC infer-experiment.py. Each column represents read fraction in a cell. (F) Detected counts and gene numbers of MEF cells between scPolyA-seq and Smart-seq2(Chen et al. 2020 Front Genet).

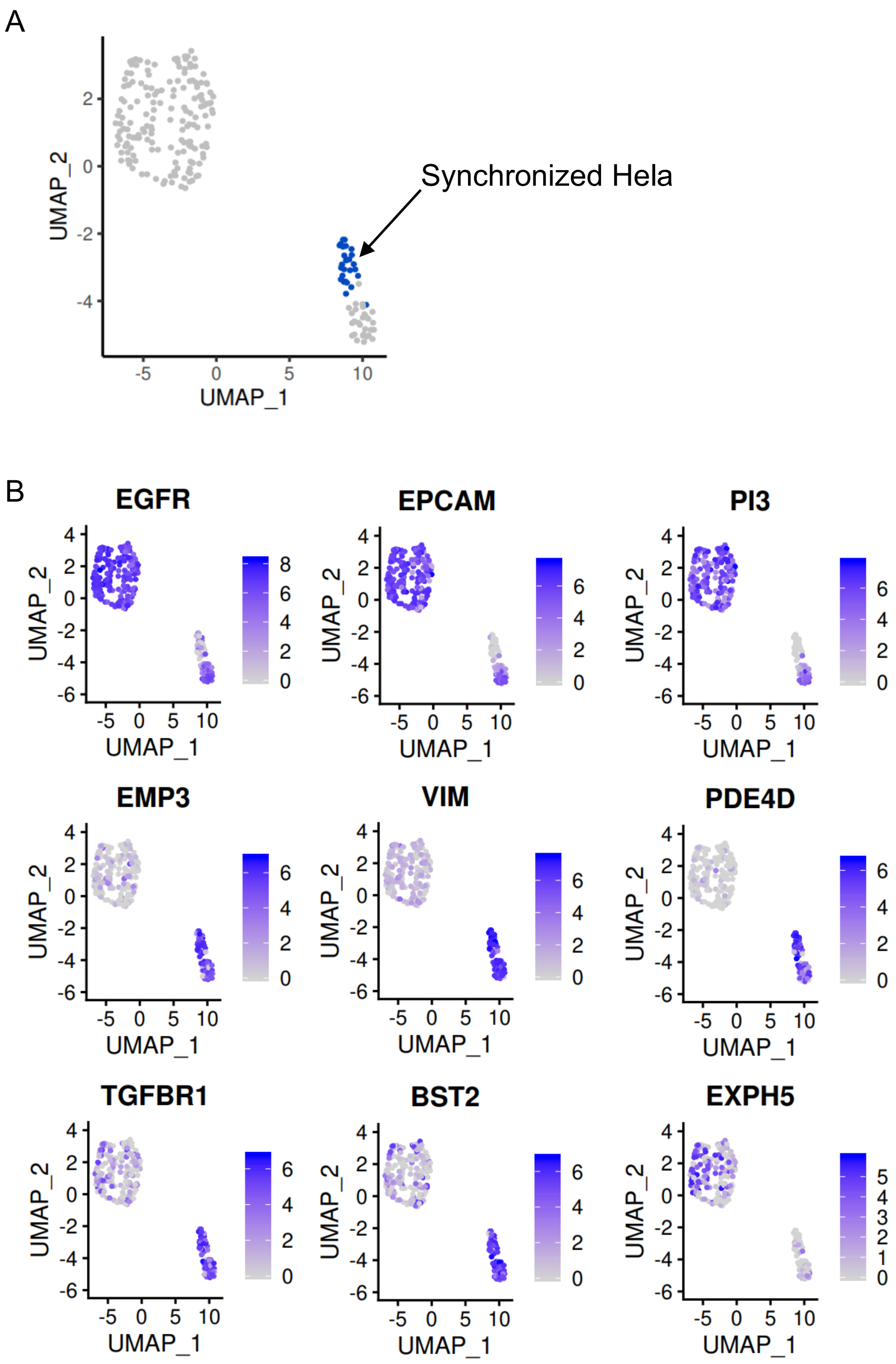


**Fig. S2. Cell clustering and cell type specific genes.** (A) UMAP plot of cells from human cell lines, with 2 sub-clusters showing up, while the top half of cluster 1 contained synchronized HeLa cells. (B) The expression levels of cell type specific genes.

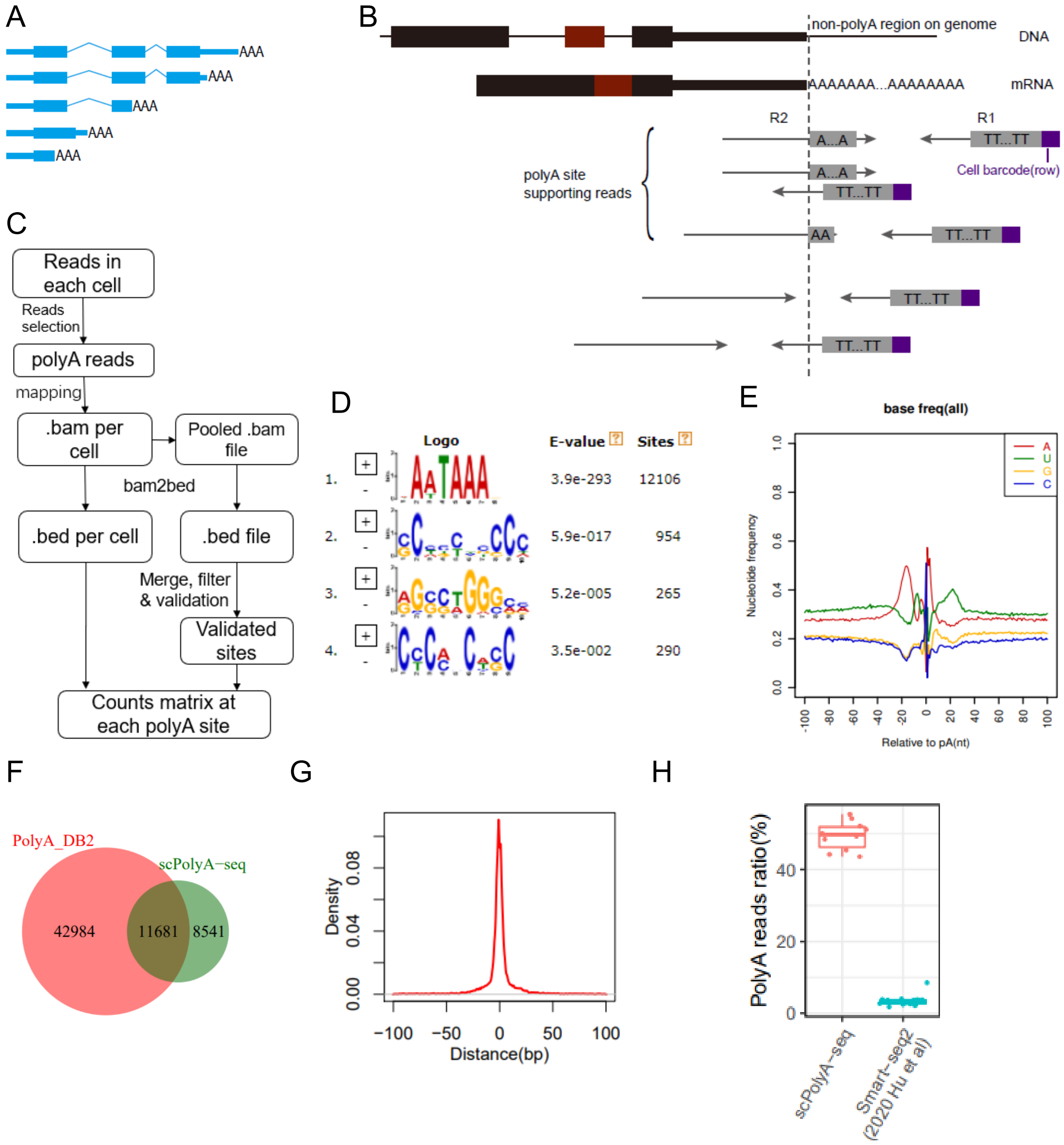


**Fig. S3. Identification of polyA sites and their features.** (A) Scheme of a gene with multiple polyA sites, either coding functional protein or not. (B) Scheme of polyA site supporting reads that used for identification of polyA site. (C) The pipeline for *de novo* polyA site identification in this study. (D) The most significantly enriched motifs near polyA sites (between -60bp and 0bp) inferred by MEME. (E) Base content frequency of 100bp upstream and downstream around each polyA site. (F) Venn diagram showed the relationship between polyA sites identified in scPolyA-seq and polyA sites in PolyA_DB2. Two polyA sites are treated as the same one if their distance <12nt. (G) Distance distribution between polyA site discovered by scPolyA-seq and the nearest polyA site in PolyA_DB2. (H) The percentage of polyA-site supporting (PASS) reads in scPolyA-seq and Smart-seq2.

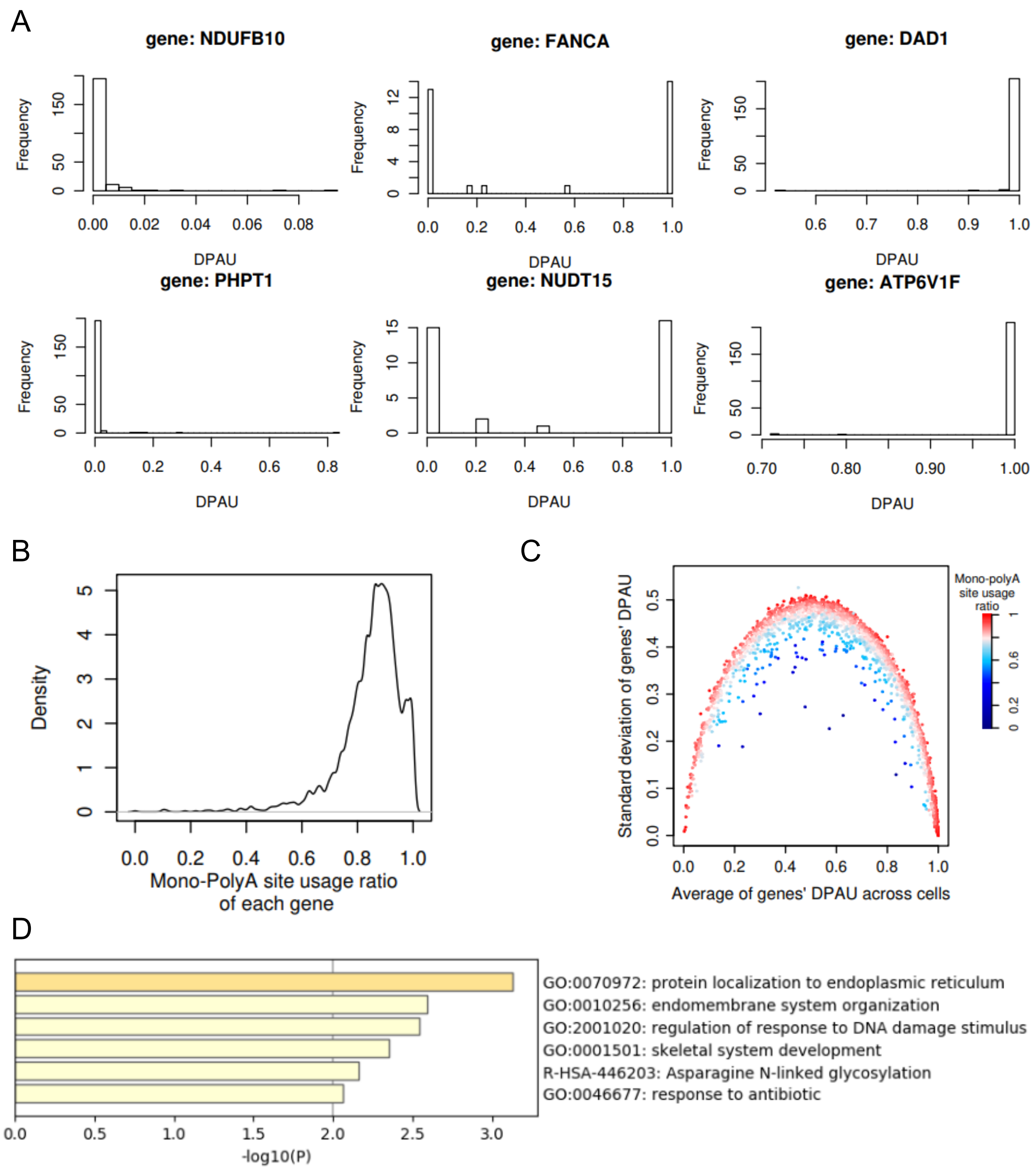


**Fig. S4. Mono-polyA site usage are prevail for genes.** (A) Histograms of DPAU of NDUFB10, PHPT1, FANCA, NUDT15, DAD1 and ATP6V1F. (B) Distribution of genes' mono-polyA site usage ratio, defined as the ratio of cells whose reads mainly at proximal or distal sites (>= 95% at either site) for each gene, shows most genes tend to favor one site in a single cell. (C) Blue points represent genes with low mono-polyA site usage ratio, which means they tend to use both sites of a gene's two poly(A) sites in one single cell, demonstrated lower variation of DPAU compared with the same level of average DPAU. (D) GO term of genes which tend to use both sites in one cell (mono-polyA site usage ratio<0.5, n=31 genes).

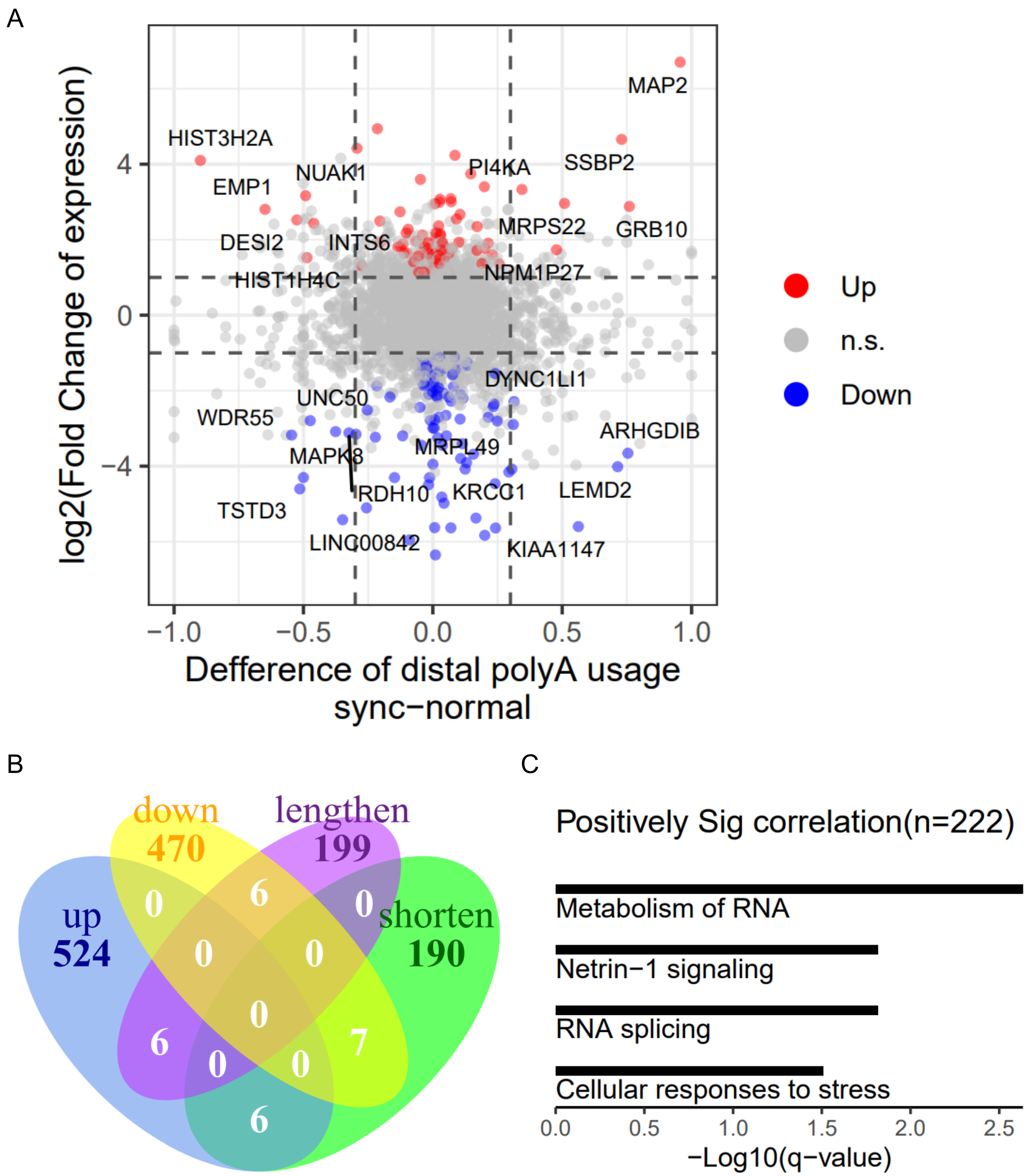


**Fig. S5. Correlation between gene expression level and APA level.** Scatterplot (A) and Venn plot (B) showed little overlaps between significant changed genes at expression level and APA level in HeLa cell before and after synchronization. (C) GO enrichment analyses of genes with significantly positive correlation between gDPAU and expression from Fig. 3A (Spearman correlation>0, FDR<0.05).

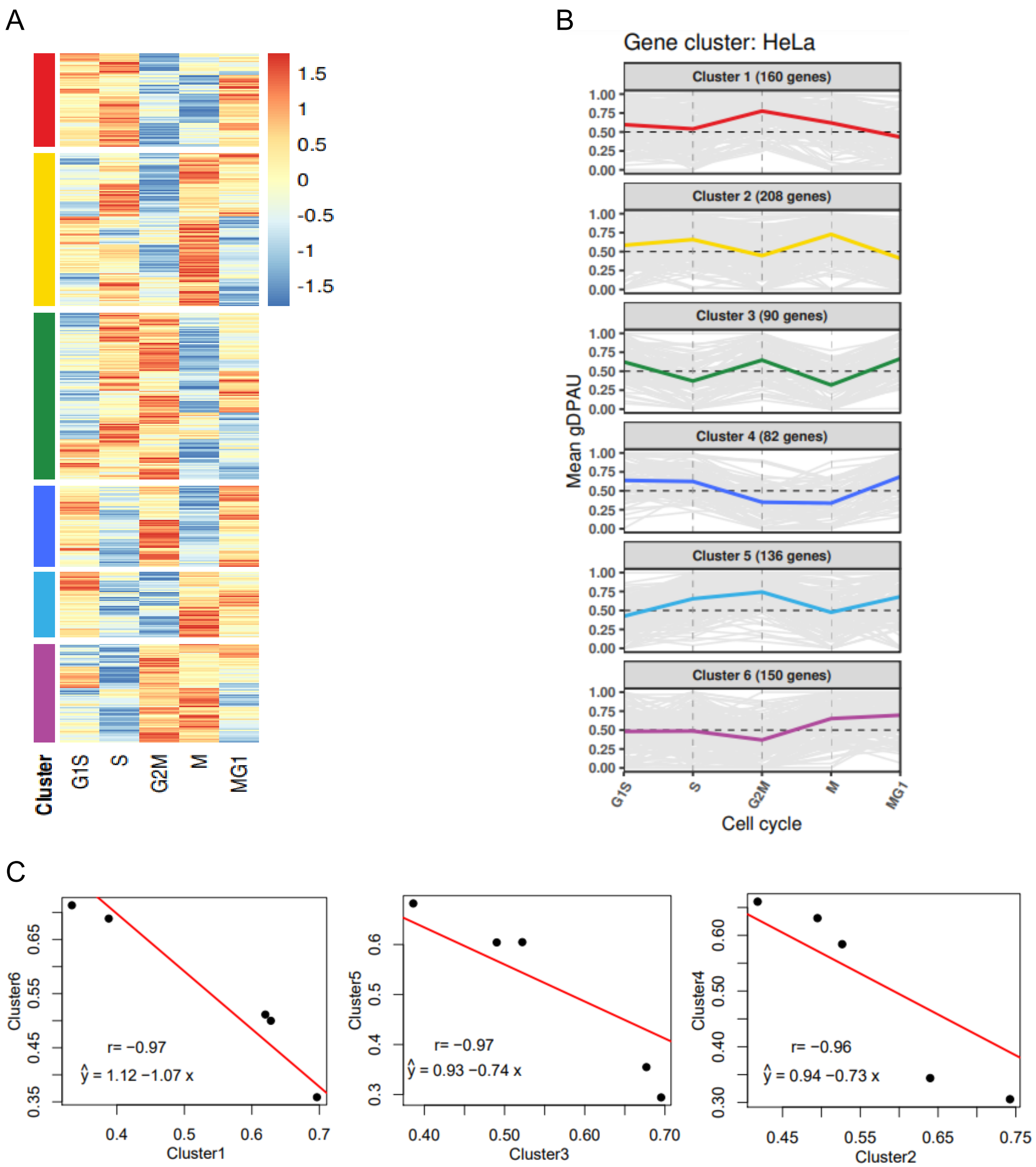


**Fig. S6. gDPAU of genes in each gene cluster along cell cycle.** (A) Heatmap of genome-wide APA dynamics along cell cycle phases in breast cancer cell line MDA-MB-468. (B) Dynamics of gDPAU of each gene at each phase for the 6 gene clusters in HeLa cells. (C) Correlation between negatively correlated cluster-pairs at APA level.

**Table S1 (separate file).** Spearman correlation between gDPAU and gene expression.

**Table S2 (separate file).** Significantly differential expressed genes between normal HeLa and synchronized HeLa by DESeq2.

**Table S3 (separate file).** PolyA site usage switched genes between synchronized HeLa and normal HeLa.

**Table S4 (separate file).** Cell cycle associated genes used to infer cell cycle phase in MDA-MB-468 cell line.

**Table S5 (separate file).** Genes showing polyA site usage switches between neighboring cell cycle phases in MDA-MB-468 can be grouped into six gene clusters.

**Table S6 (separate file).** The six gene clusters inferred using correlation of average gDPAU of each gene across five cell phases in MDA-MB-468 cells.
